# Purification and pigment analysis of diadinoxanthin-binding PSI-LHCI supercomplexes from *Euglena gracilis* strain Z

**DOI:** 10.1101/2025.06.29.662240

**Authors:** Runa Sakamoto, Koji Kato, Yoshiki Nakajima, Jian-Ren Shen, Ryo Nagao

## Abstract

*Euglena gracilis*, a phototrophic flagellate, possesses light-harvesting complexes (LHCs) with a pigment composition distinct from that of land plants and green algae, despite notable similarities in LHC polypeptide sequences to green algae. Here, we purified photosystem I-LHCI (PSI-LHCI) supercomplexes from *E. gracilis* strain Z and characterized their biochemical and spectroscopic properties. The purified complex exhibited a unique pigment profile, notably including diadinoxanthin that is typically found in red-lineage organisms, setting *E. gracilis* apart from green-lineage organisms. The absorption spectrum displayed the Qy band of chlorophyll *a* at 675□nm, while the 77-K fluorescence-emission spectrum revealed a prominent peak at 732□nm, closely resembling those in land plants. These features suggest that long-wavelength chlorophylls bound to LHCI may be evolutionarily conserved. Nevertheless, the absence of neoxanthin, lutein, and violaxanthin further differentiates the *Euglena* LHCIs from other oxyphototrophs. Together, these results illuminate the evolutionary diversification of PSI-LHCI supercomplexes and offer insights into the unique pigment-binding features of the *Euglena* light-harvesting system.

## Introduction

Photosystem I (PSI) catalyzes the light-driven electron transfer from plastocyanin or cytochrome *c*_6_ at the lumenal side of the thylakoid membrane to ferredoxin at the stromal side in oxygenic photosynthetic organisms (Brettel and Leibl 2001; Fromme et al. 2001; Golbeck 1992; Nelson and Junge 2015). In most species, PSI is associated with light-harvesting complexes (LHCs), forming PSI-LHCI supercomplexes that facilitate efficient excitation energy transfer. These pigment-protein assemblies vary considerably in both protein composition and pigment content across photosynthetic lineages (Croce and van Amerongen 2013, 2020; Hippler and Nelson 2021; Shen 2022).

*Euglena gracilis*, a phototrophic flagellate derived from secondary endosymbiosis (Novák Vanclová et al. 2020; Turmel et al. 2009), possesses an unusual carotenoid (Car) profile. Its LHCs contain diadinoxanthin and diatoxanthin—Cars typically found in red-lineage organisms such as diatoms and haptophytes (Falkowski et al. 2004)—while lacking lutein, violaxanthin, and zeaxanthin, which are common in green-lineage phototrophs such as land plants and green algae (Casper-Lindley and Björkman 1998; Cunningham Jr. and Schiff 1986; Kato et al. 2017; Nagao et al. 2021). Despite these differences, the LHC polypeptides of *E. gracilis* exhibited notable similarity to those of green algae (Houlné and Schantz 1988; Muchhal and Schwartzbach 1992), suggesting a conserved evolutionary origin. However, the PSI-LHCI supercomplex from *E. gracilis* remained unpurified, and its molecular and functional characteristics remained unexplored.

In this study, we successfully purified PSI-LHCI supercomplexes from *Euglena gracilis* strain Z and characterized their biochemical and spectroscopic properties. The complex was found to contain diadinoxanthin as a major Car component, clearly distinguishing it from PSI-LHCI supercomplexes in land plants and green algae. The absorption and 77 K fluorescence spectra revealed unique features, including a long-wavelength emission peak at 732 nm, suggesting the presence of conserved LHCI-bound chlorophyll (Chl) species. These results shed light on the evolutionary adaptations of light-harvesting systems in *E. gracilis* and expand our understanding of PSI-LHCI diversity across photosynthetic eukaryotes.

## Materials and methods

### Cell culture and thylakoid preparation

*Euglena gracilis* strain Z (hereafter referred to as *Euglena*) was cultured in a Cramer-Myers medium (Cramer and Myers 1952) supplemented with 1/1000 volume of KW21 (Daiichi Seimo) at 30 °C under continuous aeration and a photosynthetic photon flux density of 30 µmol photons m^−2^ s^−1^ (Nagao et al. 2021). Harvested cells were pelleted by centrifugation and resuspended in buffer A (20 mM Mes-NaOH (pH 6.5), 0.2 M trehalose, 5 mM CaCl_2_, and 10 mM MgCl_2_). Cell disruption was performed using glass beads under dark and cold conditions on ice, employing 19 cycles of 10□s agitation and 3□min rest (Nagao et al. 2017). Unbroken cells were removed by centrifugation at 3,000□×□g for 10□min at 4 °C. The resultant pellet was subjected to a second round of disruption under the same conditions, followed by centrifugation again at 3,000□×□g for 10□min at 4 °C. The supernatant was then centrifuged at 125,000□×□g for 20□min at 4 °C to isolate thylakoid membranes, which were resuspended in buffer A. Chl concentrations were determined in 100% methanol (Porra et al. 1989).

### Purification of PSI-LHCI supercomplexes

All purification steps were performed at 4 °C unless otherwise stated. Thylakoid membranes were solubilized in the dark with 1% (w/v) sucrose monolaurate (SM; Carbosynth) at a Chl concentration of 0.25□mg□mL^−1^ by gentle stirring for 20□min on ice. After centrifugation at 100,000□×□g for 20□min, the supernatant was applied to a HiTrap Q HP column (5□mL; Cytiva) equilibrated with buffer B (20□mM Mes-NaOH (pH 6.5), 0.2□M trehalose, and 0.03% SM). The column was washed with buffer B at a flow rate of 1.0□mL□min^−1^ until the eluate became colorless. Proteins were eluted using the following gradient: 0–5□min, 10% buffer C (buffer B with 500□mM NaCl); 5–35□min, 10–60% buffer C; 35–55□min, 60–100% buffer C; and 55–75□min, 100% buffer C (Figure 1A). The peak labeled as 1 (Figure 1) was collected and loaded onto a 10–40% linear trehalose gradient prepared in 20□mM Mes-NaOH (pH 6.5), 10□mM NaCl, and 0.03% SM. After centrifugation at 154,000□×□g for 18□h (P40ST rotor; Hitachi), the green band (indicated by a red arrow in Figure 1B) was harvested, concentrated using a 150□kDa cut-off filter (Apollo; Orbital Biosciences), and stored in liquid nitrogen.

**Figure 1.**
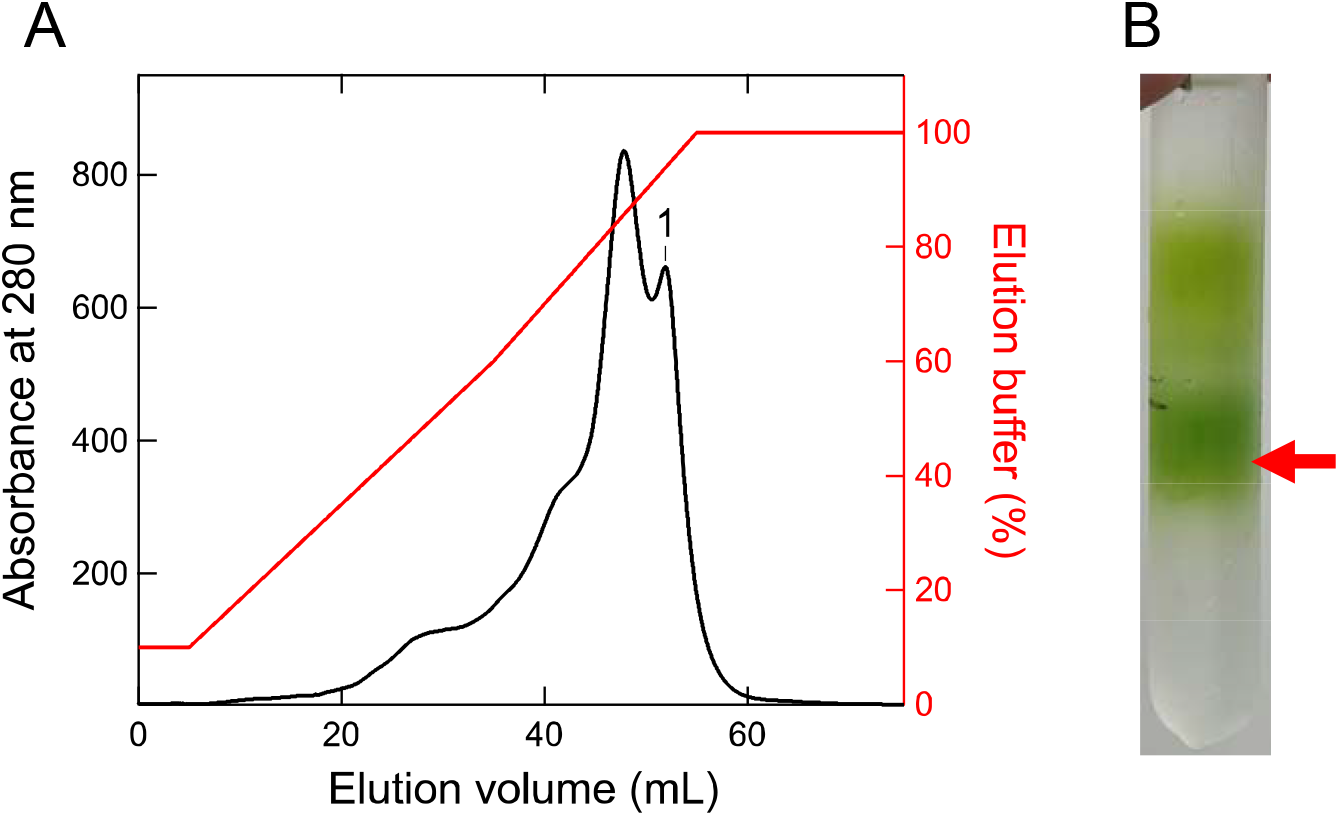
Purification of the *Euglena* PSI-LHCI. **A**, Elution profile by anion exchange chromatography. A peak labeled as 1 was collected. **B**, Trehalose density gradient centrifugation profile of the peak 1. A green fraction marked as a red arrow was collected as the *Euglena* PSI-LHCI supercomplex.

### SDS-PAGE

Protein composition was analyzed by SDS-PAGE as described by Ikeuchi and Inoue (1988). PSI-LHCI samples were solubilized in 3% lithium lauryl sulfate and 75□mM dithiothreitol at 60 °C for 10□min, and then applied to a 16% polyacrylamide gel containing 7.5□M urea. A molecular weight marker (SP-0110; APRO Science) was used. After electrophoresis, gels were stained with Coomassie Brilliant Blue R-250.

### Pigment analysis

Pigments associated with PSI-LHCI were analyzed by HPLC as described previously (Nagao et al. 2020). Pigments were extracted in 100% methanol and separated using a flow rate of 0.9□mL□min^−1^. The mobile phases consisted of solvent A (methanol:acetonitrile:0.25 M pyridine = 50:25:25 (v:v:v)) and solvent B (methanol:acetonitrile:acetone = 20:60:20 (v:v:v)) (Zapata et al. 2000). Pigments were identified by their absorption spectra and retention times (Taniguchi and Lindsey 2021; Zapata et al. 2000).

### Spectroscopic measurements

Absorption spectra were measured at room temperature using a spectrophotometer (UV-2450; Shimadzu). Fluorescence-emission spectra were recorded at 77 K using a spectrofluorometer (RF-5300PC; Shimadzu). Each of the spectra was averaged and normalized.

## Results

The PSI-LHCI supercomplexes from *E. gracilis* were purified by anion-exchange chromatography followed by trehalose density gradient centrifugation (Figure 1). The polypeptide composition of the purified complexes was examined by SDS-PAGE (Figure□2). Multiple bands were observed, including a prominent band (marked with an asterisk) corresponding to PsaA/B, as inferred from its apparent molecular weight between 45.0 and 66.4 kDa. Additional bands within the 14.3–29.0 kDa range likely represent PSI and LHCI subunits, and the overall band pattern resembled those reported for PSI-LHCI from land plants (Croce et al. 1996; Dunahay and Staehelin 1985) and green algae (Germano et al. 2002; Qin et al. 2015).

**Figure 2.**
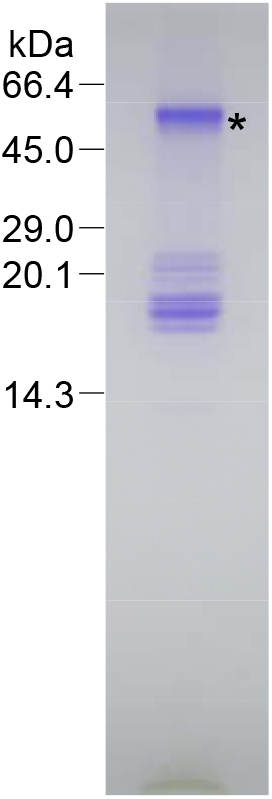
Polypeptide composition of the *Euglena* PSI-LHCI. The *Euglena* PSI-LHCI was subjected to SDS-PAGE and its protein profile showed a typical PSI protein band of PsaA/B marked as an asterisk.

The pigment composition of the *Euglena* PSI-LHCI was analyzed by HPLC (Figure□3). Four pigments—diadinoxanthin, Chl *b*, Chl *a*, and β-carotene—were detected. Diadinoxanthin appeared to be the major Car, consistent with earlier analyses of isolated *Euglena* LHCs (Cunningham Jr. and Schiff 1986). The Car composition differed markedly from that of PSI-LHCI in land plants and green algae, whose LHCs lack diadinoxanthin (Klimmek et al. 2005; Qin et al. 2015; Schmid et al. 2002; Shen 2022).

**Figure 3.**
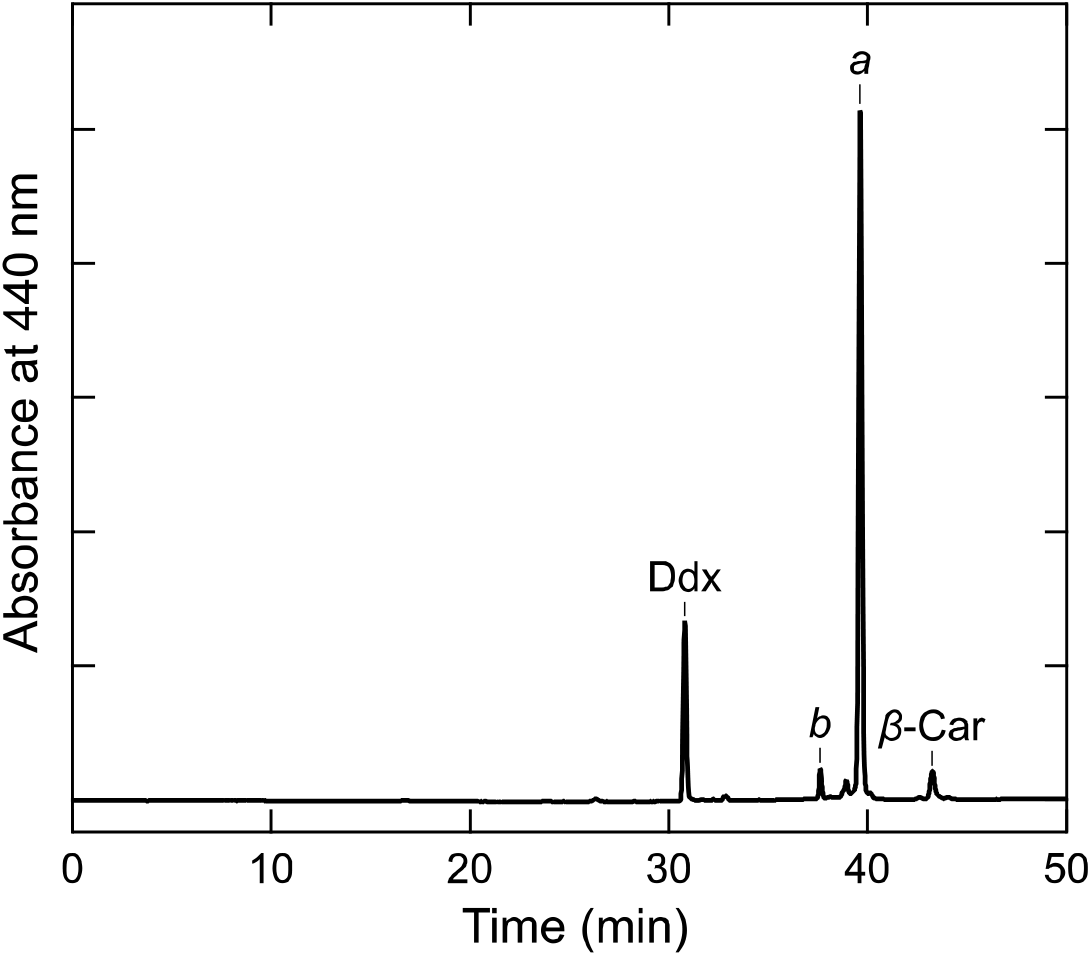
HPLC profile of pigments extracted from the *Euglena* PSI-LHCI. The HPLC chromatogram of the *Euglena* PSI-LHCI was monitored at 440 nm. Ddx, diadinoxanthin; *b*, Chl *b*; *a*, Chl *a*; β-Car, β-carotene.

The absorption spectrum of *Euglena* PSI-LHCI measured at room temperature showed the Qy band of Chl *a* at 675□nm and additional peaks attributable to Chls and Cars in the 400–500□nm region (Figure□4). Compared with PSI-LHCI from *Spinacia oleracea* (Qin et al. 2006) and *Chlamydomonas reinhardtii* (Drop et al. 2011), the *Euglena* complex exhibited relatively lower absorbance in the 450–500 nm range, likely reflecting differences in Car composition, particularly the absence of lutein and neoxanthin. Furthermore, the Qy peak of Chl *a* in the *Euglena* PSI-LHCI appeared at a shorter wavelength (675□nm) than those in *S. oleracea* and *C. reinhardtii* (679□nm) (Drop et al. 2011; Qin et al. 2006), implying differences in Chl composition and binding properties.

**Figure 4.**
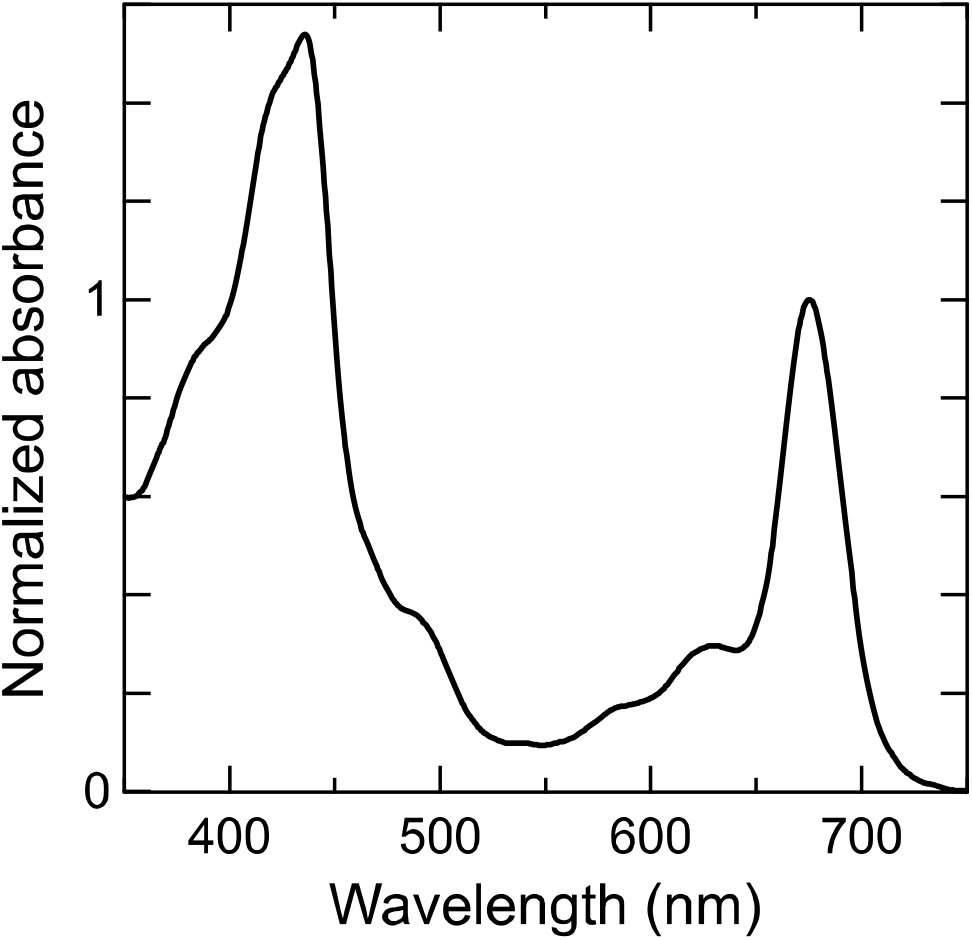
Absorption spectrum of the *Euglena* PSI-LHCI. The absorption spectra of the *Euglena* PSI-LHCI were measured at room temperature, and then averaged and normalized to the intensity of the Qy band of Chl *a*.

The 77-K fluorescence-emission spectrum of *Euglena* PSI-LHCI displayed a maximum peak at 732 nm and a broad shoulder around 677 nm (Figure□5). The 732-nm peak closely matches that observed in PSI-LHCI from land plants (Drop et al. 2011; Qin et al. 2006), but differs from the 714-nm peak characteristic of *C. reinhardtii* (Le Quiniou et al. 2015; Turmel et al. 2009). As the long-wavelength fluorescence near 730□nm originates from LHCI-bound Chls in land plants (Croce et al. 1998; Lam et al. 1984; Pålsson et al. 1995; Qin et al. 2006), it is plausible that the 732-nm emission in *Euglena* also arises from LHCI-associated Chls. In contrast, the 677-nm emission may originate from higher-energy Chls within LHCIs and/or dissociated LHCIs, consistent with the fluorescence maximum of isolated LHCIs from *S. oleracea*, which peaked at 675–680 (Lam et al. 1984; Qin et al. 2006).

**Figure 5.**
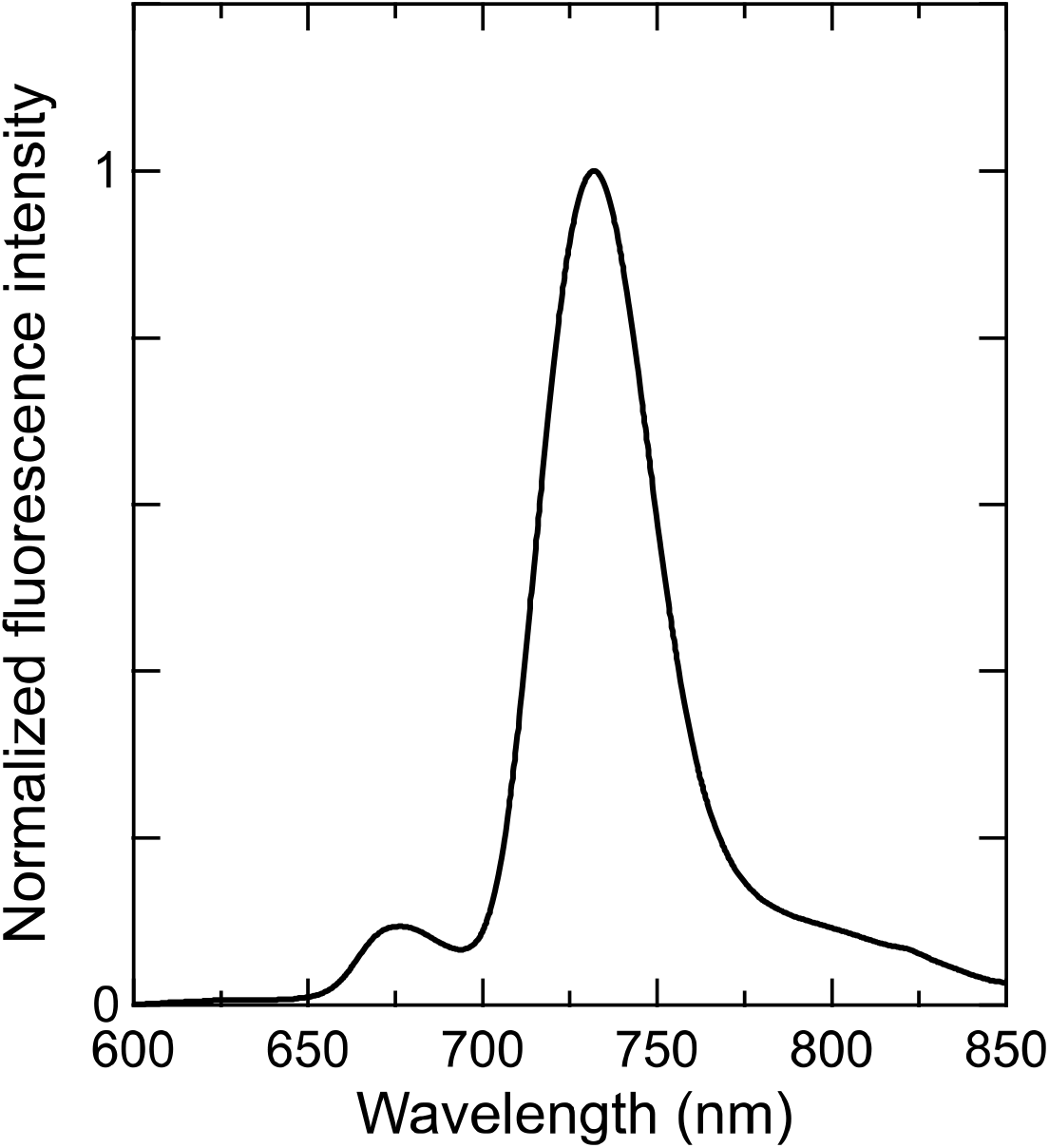
Fluorescence emission spectrum of the *Euglena* PSI-LHCI. The fluorescence emission spectra of the *Euglena* PSI-LHCI were measured at 77 K upon excitation at 430 nm, and then averaged and normalized to the maximum-peak intensity.

## Discussion

In this study, we encountered considerable difficulty in isolating thylakoid membranes from *Euglena gracilis*. Conventionally, thylakoid membranes are obtained by pelleting the supernatant following low-speed centrifugation after glass bead disruption (see Materials and Methods). However, in *Euglena*, although thylakoid membranes were successfully solubilized, the extracted protein complexes consistently aggregated during anion-exchange chromatography and trehalose density gradient centrifugation. In contrast, reproducible isolation of PSI-LHCI was achieved when the initial low-speed pellet was subjected to an additional round of glass bead disruption. Although the mechanistic basis remains unclear, *Euglena* cultured under our conditions required this modified approach for successful thylakoid membrane preparation.

The Car composition of the *Euglena* PSI-LHCI was markedly distinct from that of land plants and green algae (Figure 2). In particular, the presence of diadinoxanthin is notable, as this Car is typically found in red-lineage organisms. Conversely, neoxanthin, lutein, and violaxanthin—dominant in the LHCs of green-lineage organisms—were absent. Given that the LHC proteins of *Euglena* share sequence similarity with those of green algae (Houlné and Schantz 1988; Muchhal and Schwartzbach 1992) and are classified within the LHC protein superfamily (Engelken et al. 2010; Sturm et al. 2013), a degree of structural conservation is expected. Nevertheless, the divergent Car profile raises important questions about the mechanisms underlying Car binding specificity in *Euglena* LHCs.

The 77-K fluorescence-emission spectrum of *Euglena* PSI-LHCI exhibited a maximum at 732 nm (Figure□5), closely resembling that of *S. oleracea* (Qin et al. 2006) yet distinct from the 714-nm peak characteristic of *C. reinhardtii* (Le Quiniou et al. 2015; Turmel et al. 2009). In other plant species, major long-wavelength fluorescence peaks near 750 nm have also been reported (Li et al. 2024). Structural analyses have attributed such long-wavelength emission to Chls bound within LHCI (Li et al. 2024), raising the possibility that the 732-nm-emitting Chls in *Euglena* are likewise associated with LHCIs.

In summary, the presence of diadinoxanthin and the absence of canonical green-lineage Cars underscores the distinct pigment composition of the *Euglena* PSI-LHCI, highlighting its divergence from typical land plant and green algae. Nevertheless, the 732-nm fluorescence emission implies a degree of functional conservation with LHCI in land plants. Despite the similarity in LHC protein sequences to green algae, the unique Car profile in *Euglena* suggests independent adaptations in pigment-binding properties. These findings broaden our understanding of PSI-LHCI diversity and offer new perspectives on the structural and pigment-binding variations in photosynthetic light-harvesting systems.

## Abbreviations

Car: carotenoid
Chl: chlorophyll
LHC: light-harvesting complex
PSI: photosystem I
SM: sucrose monolaurate.

## Acknowledgements

We thank Kumiyo Kato and Satoko Kakiuchi for helpful assistance in this study. The cells of *E. gracilis* strain Z were given by Prof. Takahiro Ishikawa, Shimane University, Japan. This work was supported by JSPS KAKENHI grant Nos. JP23K14211 (Y.N.), JP22H04916 (J.-R.S.), and JP23H02423 (R.N.) and Takeda Science Foundation (K.K.).

## Author Contributions

R.N. conceived the project; R.S. performed isolation and characterization of the PSI-LHCI supercomplexes; K.K, Y.N., J.-R.S., and R.N. provided laboratory resources and supervised experimental work; R.N. drafted the original manuscript; J.-R.S. modified the manuscript; and R.N. wrote the final manuscript, and all authors joined the discussion of the results.

## Declaration of competing interest

The authors declare no conflict of interest.

## Notes

### Competing Interest Statement

The authors have declared no competing interest.

## References

Brettel K, Leibl W (2001) Electron transfer in photosystem I. Biochim Biophys Acta, Bioenerg 1507: 100–114.

Casper-Lindley C, Björkman O (1998) Fluorescence quenching in four unicellular algae with different light-harvesting and xanthophyll-cycle pigments. Photosynth Res 56: 277–289.

Cramer M, Myers J (1952) Growth and photosynthetic characteristics of Euglena gracilis. Arch Mikrobiol 17: 384–402.

Croce R, van Amerongen H (2013) Light-harvesting in photosystem I. Photosynth Res 116: 153–166.

Croce R, van Amerongen H (2020) Light harvesting in oxygenic photosynthesis: structural biology meets spectroscopy. Science 369.

Croce R, Zucchelli G, Garlaschi FM, Bassi R, Jennings RC (1996) Excited state equilibration in the photosystem I-light-harvesting I complex: P700 is almost isoenergetic with its antenna. Biochemistry 35: 8572–8579.

Croce R, Zucchelli G, Garlaschi FM, Jennings RC (1998) A thermal broadening study of the antenna chlorophylls in PSI-200, LHCI, and PSI core. Biochemistry 37: 17355–17360.

Cunningham Jr. FX, Schiff JA (1986) Chlorophyll-protein complexes from Euglena gracilis and mutants deficient in chlorophyll b: I. pigment composition. Plant Physiol 80: 223–230.

Drop B, Webber-Birungi M, Fusetti F, Kouřil R, Redding KE, Boekema EJ, Croce R (2011) Photosystem I of Chlamydomonas reinhardtii contains nine light-harvesting complexes (Lhca) located on one side of the core. J Biol Chem 286: 44878–44887.

Dunahay TG, Staehelin LA (1985) Isolation of photosystem I complexes from octyl glucoside/sodium dodecyl sulfate solubilized spinach thylakoids : characterization and reconstitution into liposomes. Plant Physiol 78: 606–613.

Engelken J, Brinkmann H, Adamska I (2010) Taxonomic distribution and origins of the extended LHC (light-harvesting complex) antenna protein superfamily. BMC Evol Biol 10: 233.

Falkowski PG, Katz ME, Knoll AH, Quigg A, Raven JA, Schofield O, Taylor FJR (2004) The evolution of modern eukaryotic phytoplankton. Science 305: 354–360.

Fromme P, Jordan P, Krauß N (2001) Structure of photosystem I. Biochim Biophys Acta, Bioenerg 1507: 5–31.

Germano M, Yakushevska AE, Keegstra W, van Gorkom HJ, Dekker JP, Boekema EJ (2002) Supramolecular organization of photosystem I and light-harvesting complex I in Chlamydomonas reinhardtii. FEBS Lett 525: 121–125.

Golbeck JH (1992) Structure and function of photosystem I. Ann Rev Plant Physiol Plant Mol Biol 43: 293–324.

Hippler M, Nelson N (2021) The plasticity of photosystem I. Plant Cell Physiol 62: 1073–1081.

Houlné G, Schantz R (1988) Characterization of cDNA sequences for LHCI apoproteins in Euglena gracilis: the mRNA encodes a large precursor containing several consecutive divergent polypeptides. Mol Gen Genet 213: 479–486.

Ikeuchi M, Inoue Y (1988) A new photosystem II reaction center component (4.8 kDa protein) encoded by chloroplast genome. FEBS Lett 241: 99–104.

Kato S, Soshino M, Takaichi S, Ishikawa T, Nagata N, Asahina M, Shinomura T (2017) Suppression of the phytoene synthase gene (EgcrtB) alters carotenoid content and intracellular structure of Euglena gracilis. BMC Plant Biol 17: 125.

Klimmek F, Ganeteg U, Ihalainen JA, van Roon H, Jensen PE, Scheller HV, Dekker JP, Jansson S (2005) Structure of the higher plant light harvesting complex I: in vivo characterization and structural interdependence of the Lhca proteins. Biochemistry 44: 3065–3073.

Lam E, Oritz W, Mayfield S, Malkin R (1984) Isolation and characterization of a light-harvesting chlorophyll a/b protein complex associated with photosystem I. Plant Physiol 74: 650–655.

Le Quiniou C, Tian L, Drop B, Wientjes E, van Stokkum IHM, van Oort B, Croce R (2015) PSI-LHCI of Chlamydomonas reinhardtii: increasing the absorption cross section without losing efficiency. Biochim Biophys Acta, Bioenerg 1847: 458–467.

Li X, Huang G, Zhu L, Hao C, Sui S-F, Qin X (2024) Structure of the red-shifted Fittonia albivenis photosystem I. Nat Commun 15: 6325.

Muchhal US, Schwartzbach SD (1992) Characterization of a Euglena gene encoding a polyprotein precursor to the light-harvesting chlorophyll a/b-binding protein of photosystem II. Plant Mol Biol 18: 287–299.

Nagao R, Ueno Y, Akimoto S, Shen J-R (2020) Effects of CO2 and temperature on photosynthetic performance in the diatom Chaetoceros gracilis. Photosynth Res 146: 189–195.

Nagao R, Yamaguchi M, Nakamura S, Ueoka-Nakanishi H, Noguchi T (2017) Genetically introduced hydrogen bond interactions reveal an asymmetric charge distribution on the radical cation of the special-pair chlorophyll P680. J Biol Chem 292: 7474–7486.

Nagao R, Yokono M, Kato K-H, Ueno Y, Shen J-R, Akimoto S (2021) High-light modification of excitation-energy-relaxation processes in the green flagellate Euglena gracilis. Photosynth Res 149: 303–311.

Nelson N, Junge W (2015) Structure and energy transfer in photosystems of oxygenic photosynthesis. Annu Rev Biochem 84: 659–683.

Novák Vanclová AMG, Zoltner M, Kelly S, Soukal P, Záhonova K, Füssy Z, Ebenezer TE, Lacová Dobáková E, Eliáš M, Lukeš J, Field MC, Hampl V (2020) Metabolic quirks and the colourful history of the Euglena gracilis secondary plastid. New Phytol 225: 1578–1592.

Pålsson L-O, Tjus SE, Andersson B, Gillbro T (1995) Ultrafast energy-transfer dynamics resolved in isolated spinach light-harvesting complex I and the LCHI-730 subpopulation. Biochim Biophys Acta, Bioenerg 1230: 1–9.

Porra RJ, Thompson WA, Kriedemann PE (1989) Determination of accurate extinction coefficients and simultaneous equations for assaying chlorophylls a and b extracted with four different solvents: verification of the concentration of chlorophyll standards by atomic absorption spectroscopy. Biochim Biophys Acta, Bioenerg 975: 384–394.

Qin X, Wang K, Chen X, Qu Y, Li L, Kuang T (2006) Rapid purification of photosystem I chlorophyll-binding proteins by differential centrifugation and vertical rotor. Photosynth Res 90: 195–204.

Qin X, Wang W, Chang L, Chen J, Wang P, Zhang J, He Y, Kuang T, Shen J-R (2015) Isolation and characterization of a PSI-LHCI super-complex and its sub-complexes from a siphonaceous marine green alga, Bryopsis Corticulans. Photosynth Res 123: 61–76.

Schmid VHR, Potthast S, Wiener M, Bergauer V, Paulsen H, Storf S (2002) Pigment binding of photosystem I light-harvesting proteins. J Biol Chem 277:37307–37314.

Shen J-R (2022) tStructure, function, and variations of the photosystem I-antenna supercomplex from different photosynthetic organisms. In: Harris JR, Marles-Wright J (eds) Macromolecular Protein Complexes IV. Subcellular Biochemistry,vol 99. Springer, Cham., pp 351–377

Sturm S, Engelken J, Gruber A, Vugrinec S, Kroth PG, Adamska I, Lavaud J (2013) A novel type of light-harvesting antenna protein of red algal origin in algae with secondary plastids. BMC Evol Biol 13: 159.

Taniguchi M, Lindsey JS (2021) Absorption and fluorescence spectral database of chlorophylls and analogues. Photochem Photobiol 97: 136–165.

Turmel M, Gagnon M-C, O’Kelly CJ, Otis C, Lemieux C (2009) The chloroplast genomes of the green algae Pyramimonas, Monomastix, and Pycnococcus shed new light on the evolutionary history of prasinophytes and the origin of the secondary chloroplasts of euglenids. Mol Biol Evol 26: 631–648.

Zapata M, Rodríguez F, Garrido JL (2000) Separation of chlorophylls and carotenoids from marine phytoplankton: a new HPLC method using a reversed phase C_8_ column and pyridine-containing mobile phases. Mar Ecol Prog Ser 195: 29–45.

